# A framework for deriving analytic long-term behavior of biochemical reaction networks

**DOI:** 10.1101/2022.12.07.518183

**Authors:** Bryan S. Hernandez, Patrick Vincent N. Lubenia, Matthew D. Johnston, Jae Kyoung Kim

## Abstract

The long-term behaviors of biochemical systems are described by their steady states. Deriving these states directly for complex networks arising from real-world applications, however, is often challenging. Recent work has consequently focused on network-based approaches. Specifically, biochemical reaction networks are transformed into weakly reversible and deficiency zero networks, which allows the derivation of their analytic steady states. Identifying this transformation, however, can be challenging for large and complex networks. In this paper, we address this difficulty by breaking the complex network into smaller independent subnetworks and then transforming the subnetworks to derive the analytic steady states of each subnetwork. We show that stitching these solutions together leads to the the analytic steady states of the original network. To facilitate this process, we develop a user-friendly and publicly available package, COMPILES (COMPutIng anaLytic stEady States). With COMPILES, we can easily test the presence of bistability of a CRISPRi toggle switch model, which was previously investigated via tremendous number of numerical simulations and within a limited range of parameters. Furthermore, COMPILES can be used to identify absolute concentration robustness (ACR), the property of a system that maintains the concentration of particular species at a steady state regardless of any initial concentrations. Specifically, our approach completely identifies all the species with and without ACR in a complex insulin model. Our method provides an effective approach to analyzing and understanding complex biochemical systems.

**Author summary:** Steady states describe the long-term behaviors of biochemical systems, which are typically based on ordinary differential equations. To derive a steady state analytically, significant attention has been given in recent years to network-based approaches. While this approach allows a steady state to be derived as long as a network has a special structure, complex and large networks rarely have this structural property. We address this difficulty by breaking the network into smaller and more manageable independent subnetworks, and then use the network-based approach to derive the analytic steady state of each subnetwork. Stitching these solutions together allows us to derive the analytic steady state of the original network. To facilitate this process, we develop a user-friendly and publicly available package, COMPILES. COMPILES identifies critical biochemical properties such as the presence of bistability in a genetic toggle switch model and absolute concentration robustness in a complex insulin signaling pathway model.

## Introduction

Chemical reaction networks (CRNs) are fundamental in disciplines such as systems biology [1–4] and industrial chemistry [5]. The dynamical behavior of such networks is frequently modeled using a system of ordinary differential equations (ODEs) based on mass-action kinetics [6]. It is particularly important to determine properties of the steady states of ODEs because they describe long-term behaviors of the CRNs. In practice, however, this is made challenging by the high-dimensionality, nonlinearities, and typically unknown parameter values of such systems. These factors make traditional tools such as numerical studies and bifurcation analysis impractical or impossible.

Significant attention has consequently been given in recent years to CRNs with special structures in their underlying interaction networks, such as being weakly reversible (WR) and deficiency zero (DZ) [7]. A CRN is WR if the network is the union of reaction cycles, while the deficiency is a nonnegative integer which measures the dependency of the reactions. It is known that, regardless of the dimension of the system or the rate parameter values, the mass-action system associated with a WR and DZ network admits a unique locally stable steady state within each positive stoichiometric class [8–10]. The steady state set is furthermore known to have a monomial parametrization, i.e., a monomial with free parameters as variables, which can be constructed directly from the reaction graph [11–13]. The construction of such parametrizations has been vital in the study of absolute concentration robustness, which is the capacity for a system to have a species whose steady state value is robust to changes in initial conditions [14–16], and multistationarity, which is the capacity of a network to have multiple stoichiometrically-compatible steady states [17,18].

In general, however, CRNs arising in applications are rarely WR or DZ. Hence, the method of *network translation*, which could transform such CRNs into WR and DZ networks while preserving their dynamics, has been developed [13, 16, 19–21]. It has also been applied to stochastic reaction networks [22]. In practice, however, the process of network translation can be challenging for two reasons: (a) it can be difficult to find a WR and DZ network translation; (b) even after the network translation, the two desired structures are rarely satisfied, in particular for large and complex networks [23, 24].

In this paper, we address these challenges by breaking the process of network translation and computation of steady-state parametrization into smaller and more manageable pieces. We start by decomposing the CRNs into stoichiometrically independent subnetworks [25, 26]. We then translate the subnetworks into WR and DZ networks by shifting reactions, i.e., swapping reactions for ones with the same stoichiometric differences while keeping the original reaction rates. If successful, we derive the steady states of each translated subnetwork independently. The steady state of the original CRN can then be derived by stitching together the steady states of the subnetworks. To facilitate this process, we have also developed a computational package called COMPILES (COMPutIng anaLytic stEady States). We demonstrate the utility of our approach on examples drawn from a CRISPRi toggle switch model [27] and a metabolic insulin signaling model [28]. In particular, we identify their critical properties such as bistability and absolute concentration robustness without an enormous number of numerical calculations, unlike previous studies.

## Results

### Derivation of analytic steady states via network decomposition and network translation

To illustrate the approach of deriving analytic steady states via network decomposition and network translation, we consider a simple CRN 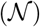 (Fig. 1a upper left). This network is not WR because the reaction *B* + *C* → *C* is not contained in a cycle. Furthermore, it is not DZ because it has six nodes (*n* = 6), two connected components (ℓ = 2), and the rank of its stoichiometric matrix is three (*s* = 3) (lower left). This gives a deficiency of *δ* = *n* – ℓ – *s* = 6 – 2 – 3 = 1 ≠ 0 (see the Methods Section for details of the deficiency). We decompose this network into two *independent subnetworks* as shown in Fig. 1a (upper right) (see the Methods Section for details on independent subnetworks). Specifically, we group the reactions of the network in such a way that the rank of the stoichiometric matrix of the original network (lower left) equals the sum of the ranks of the stoichiometric matrices of the resulting subnetworks (lower right).

**Fig 1.**
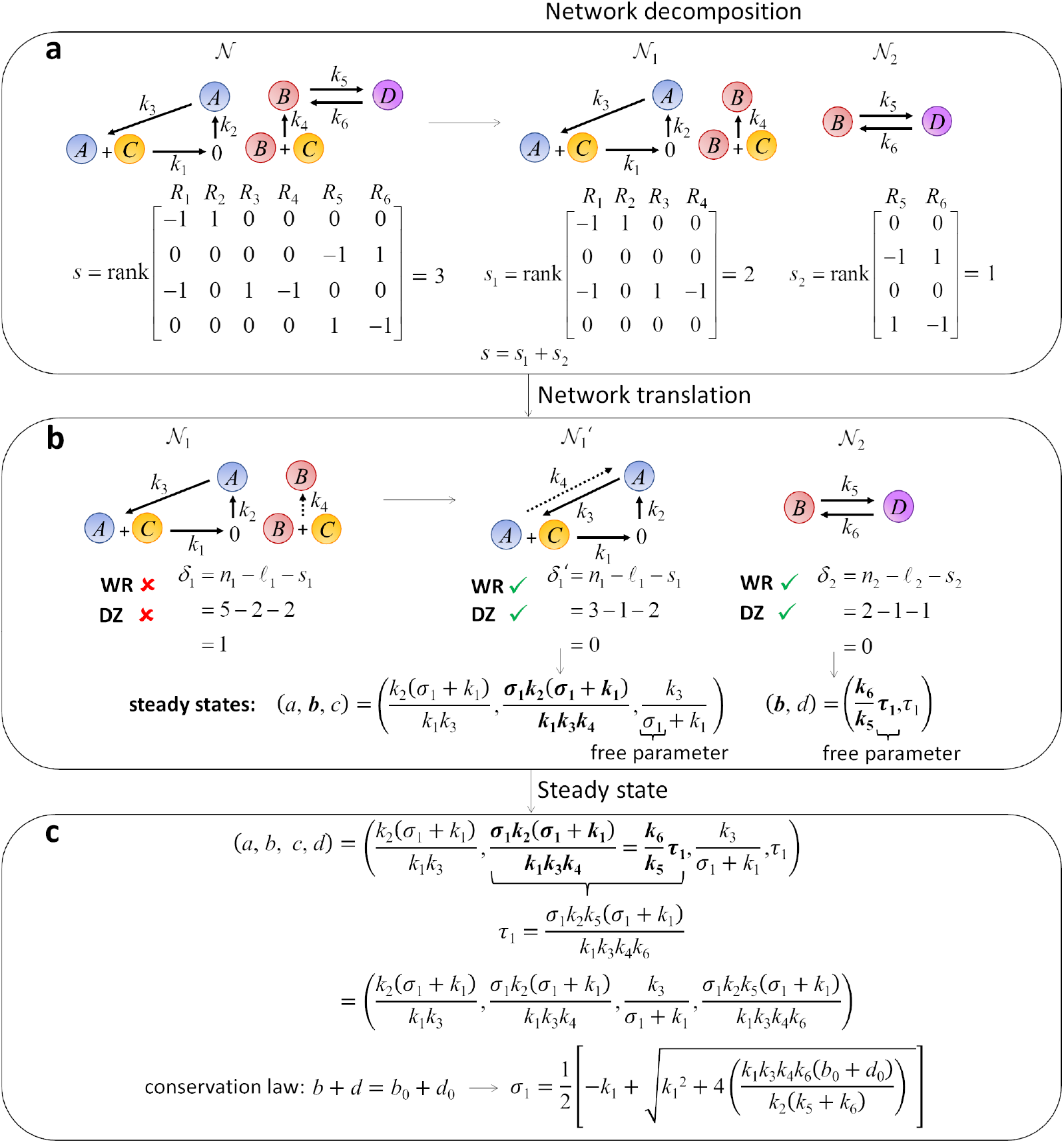
Derivation of analytic steady state via network decomposition. **a** The CRN 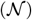 is decomposed into independent subnetworks (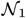 and 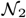) in such a way that the rank of the original matrix (*s*) is equal to the sum of the ranks of the stoichiometric matrices of subnetworks (*s*_1_ + *s*_2_). **b** Since the subnetwork 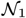 is not WR and DZ, network translation is performed. The translated subnework 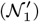 is WR and DZ while its dynamics is equivalent to the dynamics of the original subnetwork 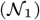. Then, the steady states of the translated subnetwork 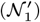 and the original subnetwork 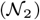 can be analytically derived since they are WR and DZ (see Fig. 2 for details). **c** Combine the steady states of subnetworks to identify the steady states of the original networks. In particular, equate the steady states of the common species of 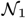 and 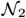 (i.e., species *B*), which eliminates the free parameter *τ*_1_. Then, by combining the steady state of every species, the steady state of the original network can be derived with one remaining free parameter *σ*_1_. This *σ*_1_ is computed from the conserved quantity in the network, which is the sum of the initial concentrations of species *B* and *D* (i.e., *b*_0_ + *d*_0_).

The second subnetwork 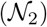 (Fig. 1b right) is a WR and DZ network. On the other hand, the first subnetwork 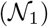 (Fig. 1b left) is neither WR nor DZ. Thus, we have to translate 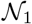 to make it WR and DZ. Specifically, 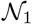 is not WR because the reaction *B* + *C* → *B* does not belong to a cycle. Thus, we replace it with *A* + *C* → *A*, having the same stoichiometry and the same reaction rate (i.e., *k*_2_*bc*) to obtain the translated subnetwork 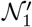 (Fig. 1b middle) (see [13] for details on network translation). We keep the original reaction rate (*k*_2_*bc*) for *A* + *C* → *A* so that the dynamics does not change. However, as the network structure changes, and in particular, as all the reactions now belong to a cycle, 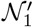 is a WR network. Furthermore, 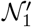 is DZ because it has three nodes (*n* = 3), one connected component (ℓ = 1), and a stoichiometric matrix of rank two (*s* = 2) so that *δ* = 3 – 1 – 2 = 0.

Since both 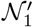 and 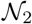 are WR and DZ, we can derive their analytic steady states. To do this, we first construct a *generalized CRN* (*GCRN*) (Fig. 2a), which is a directed graph with edges taken from the translated network (Fig. 2a (ii)) and nodes from the original network (Fig. 2a (i)) (see [12, 29] for details on GCRNs). For instance, to construct a GCRN representation for the translated network 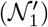, we first take all the edges of 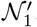. Then, we fill 0 in the tail of the edge associated with *k*_2_ (Fig. 2a (iii)) because the source node of the edge associated with *k*_2_ in the original network 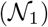 is 0 (Fig. 2a (i)). Similarly, we fill *A* in the tail of the edge associated with *k*_3_ (2a (iii)). Because the source nodes of the edges with *k*_1_ and *k*_4_ are *A* + *C* and *B* + *C* in the original network (2a (i)), two source nodes (*A* + *C* and *B* + *C*) are now placed in a single node ((2a (iii)). To separate the two source nodes, a phantom edge (i.e., an edge with a free parameter (*σ*_1_) is added (2a (iv)) (see [21] for details on phantom edges). For the phantom edge, a zero stoichiometric vector is always assigned to maintain the dynamics of the original network. This produces a GCRN representation of the translated network.

**Fig 2.**
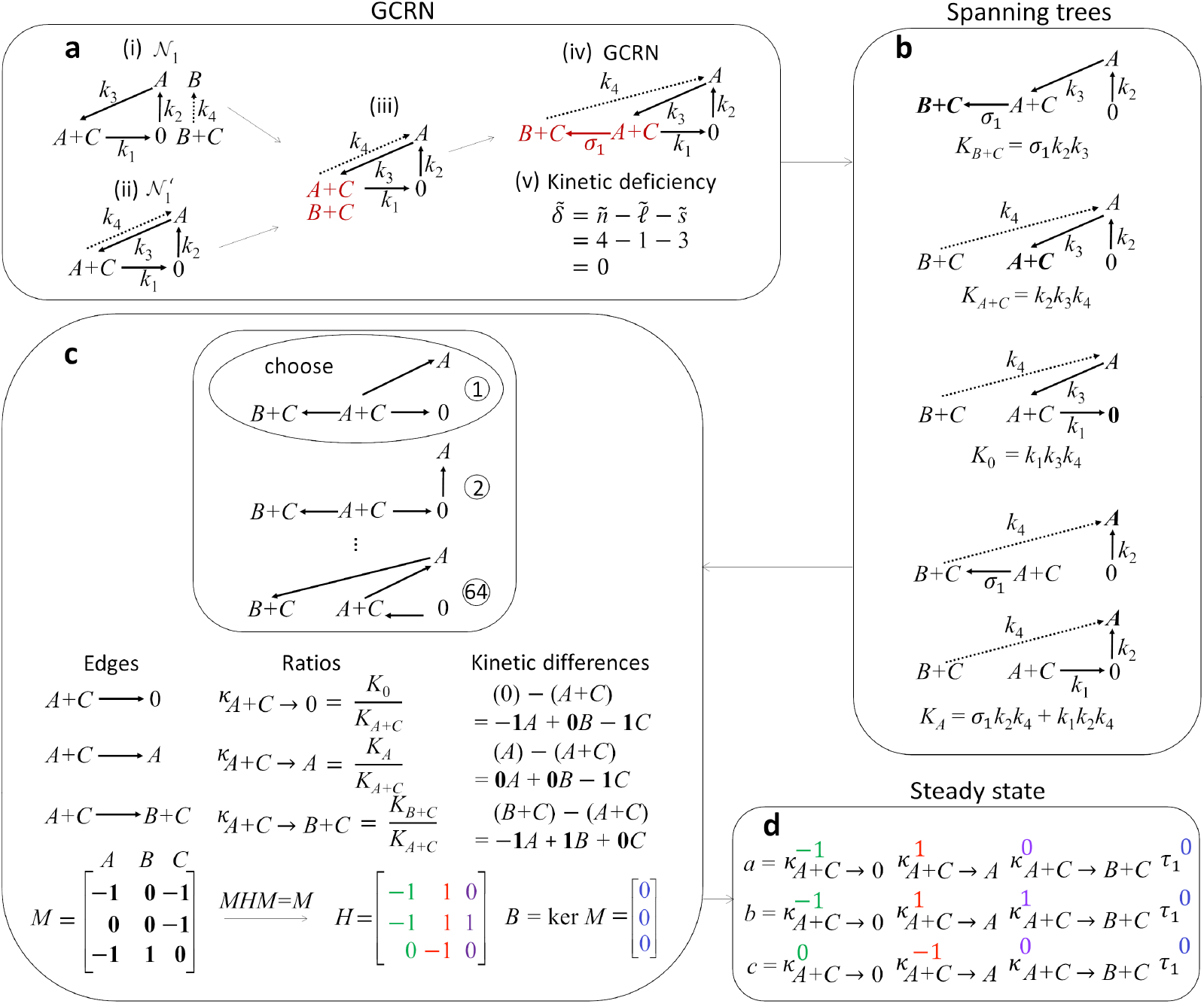
Derivation of analytic steady state via network translation. **a** The generalized CRN (GCRN) is constructed from the edges of the translated network but the nodes are the source nodes of the associated reactions in the original network. For instance, the edges associated with *k*_1_ and *k*_4_ in the translated network (ii) have corresponding source nodes *A* + *C* and *B* + *C*, respectively, in the original network (i). Since there are two source nodes (*A* + *C* and *B* + *C*) which share a single node in (iii), a phantom edge with a zero stoichiometric vector and a free parameter rate constant (*σ*_1_) is introduced (upper right). Notice that this operation maintains the dynamics of the original network. Proceed to the next step if the kinetic deficiency is zero. **b** Collect all the spanning trees of the GCRN (i.e., connected subgraphs of the GCRN without a cycle) which direct towards each node (*B* + *C*, *A* + *C*, 0, and *A*). Then multiply the rate constants associated with the edges of each spanning tree. For instance, the products of the rate constants associated with the edges of the two spanning trees towards node *A* are *σ*_1_ *k*_2_*k*_4_ and *k*_1_ *k*_2_*k*_4_ (bottom). Compute each tree constant (*K_i_*) by adding the product of the rate constants over all the spanning trees towards node (i). Hence, the tree constant *K_A_* = *σ*_1_*k*_2_*k*_4_ + *k*_1_*k*_2_*k*_4_ is obtained. **c** Choose an arbitrary tree (i.e., a connected graph without a cycle) containing all the nodes in the GCRN (top) which is not necessarily a subgraph of this GCRN. Then, for each edge (*i* → *i*′) of the chosen tree, find the ratios of the tree constants 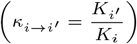 and the kinetic difference (*i*′ – *i*) (middle). For instance, the edge *A* + *C* → 0 has the ratio of the tree constants 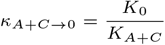 and kinetic difference (0) – (*A* + *C*) = – 1*A* + 0*B* – 1*C*. From the kinetic differences, construct the matrix *M* (bottom) by listing in each row the coefficients in the kinetic difference associated with an edge of the tree. Hence, the coefficients in the kinetic difference –1*A* + 0*B* – 1*C* are the entries of the first row of *M* (i.e., [–1, 0, –1]). Then, compute a generalized inverse *H* of *M* (i.e., MHM = M) and the kernel *B* of *M*. **d** Derive the analytic steady state of the network from the ratio of the tree constants *κ*_*i*→*i*′_ and the matrices *H* and *B*. Specifically, raise the ratios of the tree constants *κ*_*i*→*i*_′ in a component-wise manner to the entries of a row of *H*, and get their product. For instance, the ratio of the tree constants *κ*_*A*+*C*→0_, *κ*_*A*+*C*→A_, and *κ*_*A*+*C*→*B*+*C*_ are raised to the entries in the first row of *H* (– 1, 1 and 0, respectively). This gives 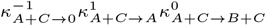. Meanwhile, one free parameter (*τ*_1_) is introduced because the number of column of *B* is one. This *τ*_1_ is raised to the entry of a row in *B*. Hence, 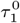 is obtained. Finally, get the product 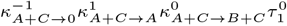, which is the steady state value of species A. The steady state values for species B and C can be computed in a similar manner.

The structure of a GCRN has its own deficiency, which is known as the *kinetic deficiency* [12, 13, 20, 29]. The formula to get the kinetic deficiency 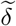 of a GCRN is the same as the one used for the original network except for the way the rank of the matrix is calculated. That is, instead of using a stoichiometric matrix, the differences of the nodes in the GCRN are used to get the rank of the matrix. The difference of the edges except for the phantom edge is the same as the stoichiometric vectors of the original translated network 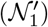, and the difference of the phantom edge ([–1,1, 0, 0]) is independent from the stoichiometric vectors of 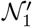. Thus, the rank of the matrix of the GCRN is increased by one compared to the 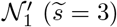. Furthermore, the GCRN has one more node (*B* + *C*) compared to 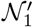, so 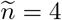. The number of connected components is not changed 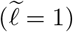. Thus, the kinetic deficiency is 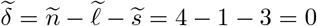 (bottom right). We proceed to the next step if the kinetic deficiency is zero; otherwise, additional conditions need to be satisfied (see [20, 21] for details).

The next step is to get all the spanning trees of the GCRN (i.e., connected subgraphs of the GCRN without a cycle) directed towards each node as shown in Fig. 2b. For instance, there is only one spanning tree directed towards *B* + *C* because there is only one connected subgraph of the GCRN without a cycle that points towards *B* + *C* (Fig. 2a (iv)). Then, to get the tree constant (*K_i_*) associated with a particular node i with only one spanning tree, we multiply the rate constants corresponding to the edges of this spanning tree. Hence, we obtain *K*_*B*+*C*_ = *σ*_1_*k*_2_*k*_4_ as the tree constant directed towards the node *B* + *C*. In general, if there is more than one spanning tree directed towards a particular node, we have to repeat the process of getting the corresponding product for each spanning tree and then get the sum of the products over all the spanning trees directed towards the particular node. Specifically, there are two spanning trees directed towards node *A* because there are two connected subgraphs of the GCRN without a cycle that point towards node *A*. Then, multiply the rate constants associated with the edges of each spanning tree. Hence, we obtain *σ*_1_*k*_2_*k*_4_ and *k*_1_*k*_2_*k*_4_ (bottom) as the products of the rate constants associated with the two spanning trees directed towards node *A*. Thus, the tree constant associated with node *A* is *K_A_* = *σ*_1_*k*_2_*k*_4_ + *k*_1_*k*_2_*k*_4_. (Note that the term “tree constant” was introduced in [13] to refer to the formulas following from the matrix tree theorem presented in [30] and adapted to chemical reaction network theory in [11].)

Next, from the GCRN (Fig. 2a (iv)), we form a tree (i.e., a connected graph without a cycle) that contains all the nodes of the GCRN. The possible trees that can be formed are given in Fig. 2c (top). Then, for each edge of the tree, find the ratio of the tree constants (center) and the kinetic difference (middle right), the ratio of the tree constants 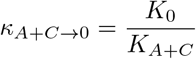, and an associated kinetic difference of (0) – (*A* + *C*) = –1*A* + 0*B* – 1*C*. After doing this for each edge of the tree, we construct the matrix *M* where each row corresponds to the vector of coefficients in the kinetic difference associated with each edge. For instance, the first row of *M* is the first kinetic difference –1*A* + 0*B* – 1*C* = [–1, 0, –1] associated with the first edge *A* + *C* → 0. In addition, the second and third rows of *M* are precisely the vectors of coefficients in the kinetic differences associated with the two remaining edges of the tree. We then compute a generalized inverse *H* of *M* (i.e., *MHM* = *M*) and the kernel *B* of *M*.

From the matrices *H* and *B* together with the ratios of the tree constants, we can derive the analytic steady state of the subnetwork 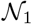 (Fig. 2d). In particular, to get the analytic steady state of the first species *A*, we raise each ratio of the tree constants associated with the three edges (i.e., *κ*_*A*+*C*→0_, *κ*_*A*+*C*→*A*_, and *κ*_*A*+*C*→*B*+*C*_) to each of the entries in the first row of *H* (i.e., − 1, 1, and 0, respectively), and then multiply them to obtain 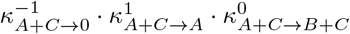. Additionally, we introduce a number of free parameters, as many as the number of columns of *B*. We have a single free parameter (*τ*_1_) because there is only one column of *B*. We raise this free parameter to the value of the first row of *B* (i.e., 0) and we get 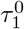. We multiply this value from the previously obtained product, which gives 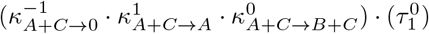. This is the value of the analytic steady state of the first species *A*. By following the same procedure, we can get the analytic steady state values of the two remaining species of the translated subnetwork 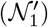. We can also get the steady state of the second subnetwork 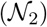 by following the same procedure (2). Note that the GCRN of 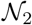 is the same as that of the original network because 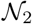 is already a WR and DZ network and thus network translation is not performed.

Then, we combine the steady states of the two subnetworks (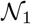 and 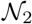) to derive the steady state of the original whole network 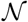. Specifically, we equate the steady state values of the common species of the two subnetworks. That is, since species *B* appears in both subnetworks, we equate the steady state values of species *B* in 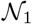 (i.e., 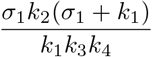) and 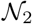 (i.e., 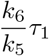) (Fig. 1b bottom). As a result, we get 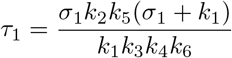. By using this, we can eliminate the free parameter (*τ*_1_) in the combined steady state (Fig. 1c upper left) and derive the analytic steady state of the whole network 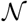 (Fig. 1c) with one free parameter *σ*_1_. This free parameter can be solved using the conservation law *b* + *d* = *b*_0_ + *d*_0_ where *b*_0_ and *d*_0_ are the initial concentrations of species *B* and *D*, respectively. That is, by substituting the steady state values of *B* and *D* to the conservation law, we can solve for the value of *σ*_1_ in terms of the conserved quantity (*b*_0_ + *d*_0_). The closed form can be used to easily identify the critical features of a steady state, such as multistability and robustness, which we will illustrate as follows.

### The analytic steady state of a simple CRISPRi toggle switch model

We now apply our method to a *CRISPRi toggle switch model* [27] with nine species (Table I in the Supplementary Information) and 14 reactions (Fig. 3a left). In the model, the deactivated mutant protein dCas9 with a single guide RNA 1 (CS complex) and dCas9 with a single guide RNA 2 (CT complex) bind to a single specific site on their target genes *H* and *G*, respectively, called *specific binding*, which forms complexes *CS* : *H* (species *P*) and *CT* : *G* (species *R*). With the model, it was shown that experimentally observed bistability of species *S* (i.e., single guide RNA 1) in response to parameter change was impossible. This leads to the identification of previously unidentified reactions of *unspecific binding* (i.e., binding to unspecific sites not matching the single guide RNA sequence, e.g., *CS* and *CT* complexes bind unspecifically to genes *G* and *H* forming complexes *CS* : *G* and *CT* : *H* as opposed to the formation of complexes *CS* : *H* and *CT* : *G* in the specific binding), which could explain the bistability in the system [27].

**Fig 3.**
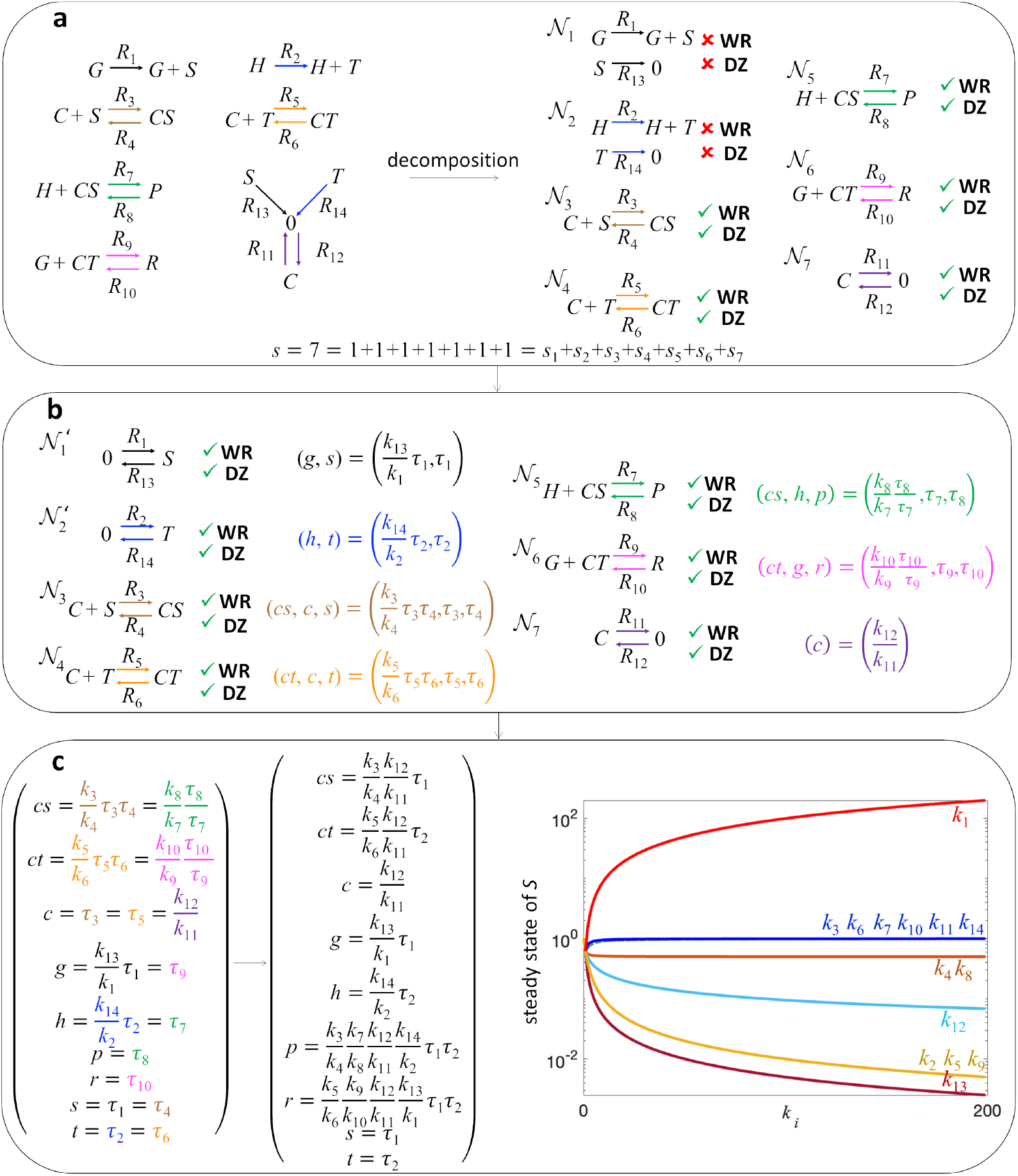
Derivation of analytic steady state of a CRISPRi toggle switch model via network decomposition and network translation. **a** A CRISPRi toggle switch network (left) that assumes dCas9 with a single guide RNA 1 (*CS* complex) and dCas9 with a single guide RNA 2 (*CT* complex) bind to a single site on their target genes *H* and *G*, respectively. The CRN has nine species (i.e., *CS, CT, C, G, H, P, R, S*, and *T*) and 14 reactions (i.e., *R*_1_,…, *R*_14_), which is decomposed into seven independent subnetworks 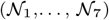 (right). **b** As the subnetworks 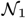 and 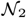 are not WR and DZ, network translation is performed. The translated subnetworks (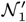 and 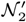) are WR and DZ while their dynamics are equivalent to the dynamics of the original subnetworks (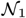 and 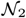, respectively). Then, the steady states of the translated subnetworks (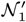 and 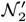) and the original subnetworks 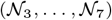 can be analytically derived as they are WR and DZ (see Fig. 2 for the outline of the steps). **c** The steady states of the subnetworks are combined by equating the steady states of the common species. For instance, the species CS is common to both subnetworks 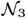 and 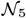 so the steady state values of CS of both subnetworks are equated to each other (left). Then after solving the steady state of the whole network, two free parameters (*τ*_1_ and *τ*_2_) are left, which can be solved in terms of the conserved quantities. This analytic steady state solution could be used to determine the behavior of the steady state concentrations of species with respect to varying rate constants. In particular, the monotonicity of the steady state concentration of species *S* over varying rate constant *k_i_*, when the rest of the rate constants are set to one, is illustrated on the right.

In a previous study, numerical simulations of the model for some parameters were used to show the absence of the bistability. In particular, in Fig. 7 of the Supplementary Information of [27], the authors considered a range of values for *k*_10_ (i.e., unbinding rate constant of the specifically bound *CT* (dCas9 with a single guide RNA 2) to *G* (Gene 1)) and performed numerical simulations to identify the effect of varying *k*_10_ on the steady state concentration of *S* (single guide RNA 1). This is time-consuming. Importantly, it was shown only for a limited choice of parameters. To circumvent this, we derive the analytic steady states of the model to show the absence of the bistability for any choice of parameters.

To do this, we decompose the model into seven independent subnetworks (Fig. 3a right). Then, for subnetworks that are not WR and DZ (i.e., 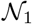 and 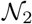), the network translation was performed to obtain WR and DZ networks. This allowed us to derive the analytic steady state of each subnetwork (Fig. 3b) using the method outlined in Fig. 2. By combining the steady states of the subnetworks, we computed the analytic form of the steady state (Fig. 3c upper left) with the values of the free parameters *τ*_1_ and *τ*_2_ using the conservation laws *h* + *p* = *h*_0_ + *p*_0_ and *g* + *r* = *g*_0_ + *r*_0_ where *h*_0_, *p*_0_, *g*_0_, *r*_0_ are the initial concentrations of species *H, P, G*, and *R*, respectively. In particular, we derived the analytic steady state of *s* as follows:

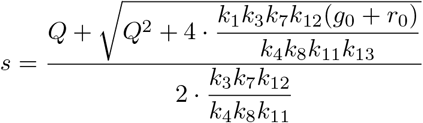

where

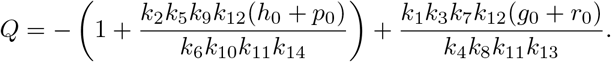

From this, we can easily show that the steady state of *S* always increases as we increase *k*_10_ because *Q* also increases as we increase *k*_10_. This confirms the absence of the bistability of *S* against *k*_10_, which was shown in a previous study under limited conditions [27]. Importantly, we can also easily observe from the formula that the steady state concentration of *S* increases monotonically over *k*_1_, *k*_6_, and *k*_14_, while monotonically decreasing over *k*_2_, *k*_5_, *k*_9_, and *k*_13_. Furthermore, the monotonicity of the steady state concentration of species S on the rest of the rate constants can be shown analytically (see the Supplementary Information for details). Hence, the bistability of S is impossible in this model for any choice of parameters. This is inconsistent with experimental observations, which indicates a missing mechanism and is consistent with the previous study [27].

### Computational package, COMPILES

Applying our method (Figs. 1 and 2) of getting the analytic steady states of a system via network decomposition and translation is challenging for complex CRNs. To resolve this, we developed a user-friendly, open-source, and publicly available computational package, COMPILES (COMPutIng anaLytic stEady States), which automatically decomposes the CRN into its finest independent decomposition and combines the solutions to the subnetworks to get the analytic steady state solution to the whole system. With this package, we were able to get the solutions to systems that involve as many as 35 reactions.

We illustrate how COMPILES derives the analytic steady state with an example of a complex CRN that describes the signaling cascades activated by insulin [28, 31] (Fig. 4a). To use the package, one simply inputs the reactions in the system’s CRN. COMPILES automatically decomposes the network into its finest independent decomposition (i.e., independent decomposition with the maximum number of subnetworks). Specifically, the package decomposes the 35 reactions of the insulin network into 10 subnetworks (Fig. 4b right). Then the steady state of each subnetwork is automatically derived using the method outlined in Figs. 1b and 2. COMPILES finally combines these subnetwork solutions to derive the analytic steady state solution for the entire network. It outputs the steady state solution with the free parameters and the conservation laws in the original network (Fig. 4b lower left).

**Fig 4.**
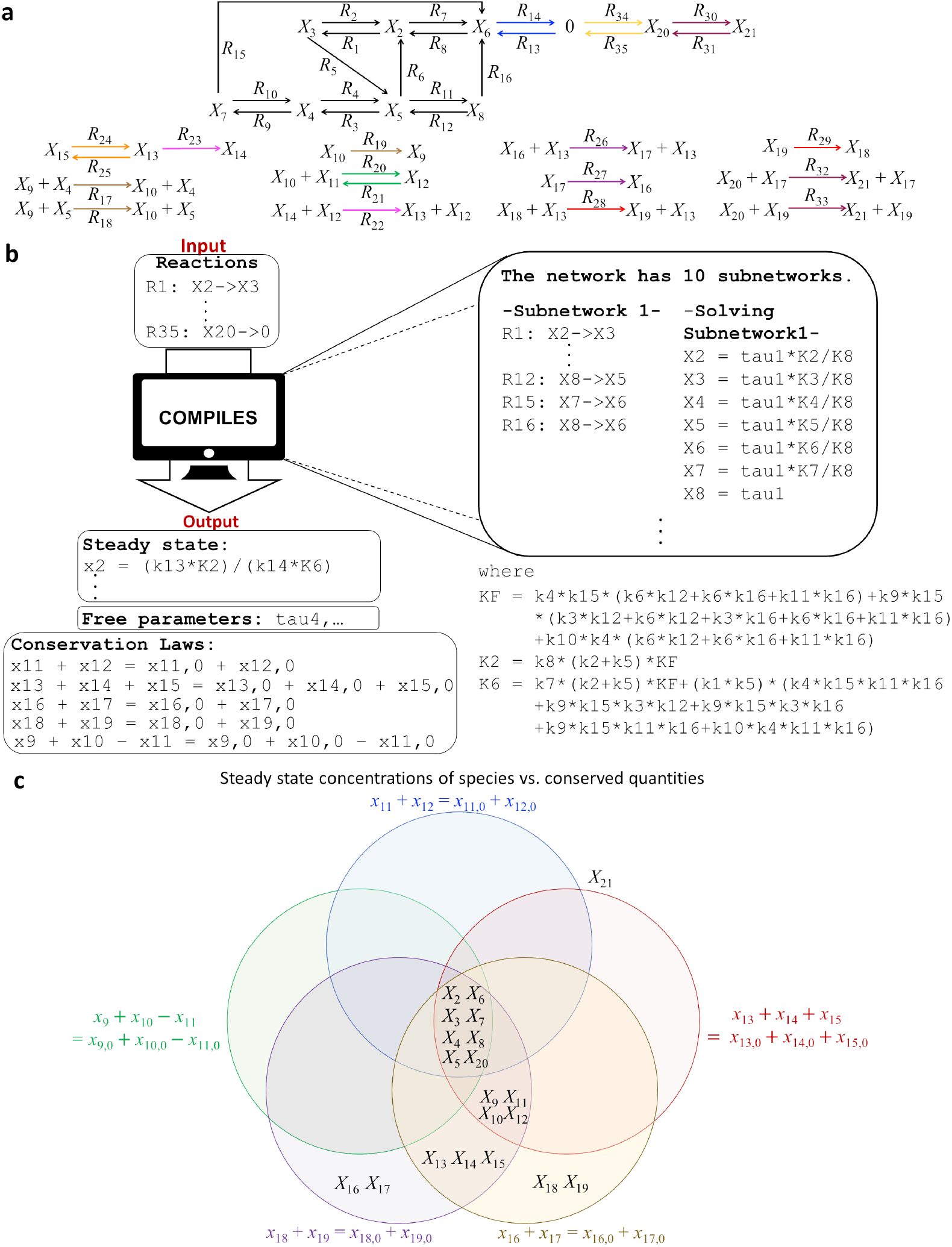
Derivation of analytic steady state of an insulin signaling network via computational package COMPILES. **a** The CRN of a mathematical model of insulin metabolic signaling pathway with 20 species (*X*_2_,…, *X*_21_) and 35 reactions (*R*_1_,…, *R*_35_). **b** A schematic diagram that describes the computational package COMPILES. As the insulin network is large, a computational package is developed to analytically derive the steady state of the network. If the users enter the reactions of the network (top left), then the package derives the steady state with the free parameters and the conservation laws of the network (bottom left). This is done by decomposing the network into independent subnetworks as outlined in Fig. 1a-b and then deriving the steady state of each subnetwork as outlined in Fig. 2 (right). Finally, the steady states of the subnetworks are combined to derive the analytic steady state of the original whole network as shown in Fig. 1c. **c** The summary of the independence of the steady state concentrations from the conserved quantities in the network when *k*_1_ = ⋯ = *k*_35_ = 1. The free parameters (i.e., *τ*_4_, *τ*_7_, *τ*_10_, *τ*_11_, and *τ*_12_) can be solved in terms of the conserved quantities. Then, a Venn diagram is used to show which among the steady state concentrations (associated with their species) are independent on specific conserved quantities. For instance, the steady state concentrations of species *X*_13_, *X*_14_, and *X*_15_ do not depend on the conserved quantities *x*_16_ + *x*_17_ = *x*_16,0_ + *x*_17,0_ (violet) and *x*_18_ + *x*_19_ = *x*_18,0_ + *x*_19,0_ (yellow); that is why these species are placed inside the violet and the yellow circles. Importantly, the steady state concentrations of species *X*_2_,…, *X*_8_, and *X*_20_ do not depend on all the initial conditions and conserved quantities (the species are placed inside all the circles), but the steady state concentration of species *X*_21_ depends on all the conserved quantities (the species is placed outside all the circles).

Using these results, we investigated the capacity for the system to exhibit *absolute concentration robustness* (*ACR*). A system is said to have ACR in a species *X* if the system has the same steady state concentration for *X* regardless of any initial concentrations [14, 15]. It basically describes the capacity to maintain the concentration of a particular species at steady state within a narrow range, regardless of the changes in the amounts of other network species that might vary due to environmental variables in the system [14]. Previously, detecting species with ACR could be done only under limited structural network conditions [14, 32]. Here, our approach allows us to detect the ACR for large networks in a manageable fashion.

To simplify the analysis of the network, we set the rate constants to 1 and the free parameters can be solved in terms of the conserved quantities. Then, we analyzed how the steady state of each species has ACR against the five conserved quantities in the system (Fig. 4c). When a species is inside a circle corresponding to a conserved quantity, it means that the steady state concentration of this species does not change even though the associated conserved quantity is varied. Interestingly, the steady states of the eight species are ACR from all five conserved quantities. In particular, one of them is *X*_20_ (intracellular GLUT4). This indicates that the long-term concentration of GLUT4 in the cell could be maintained because the system has ACR in the species *X*_20_. This allows the cell to have a certain level of glucose transporters that may be translocated to the cell surface whenever the cell needs glucose for energy metabolism. On the other hand, the system does not have ACR in species *X*_21_ (cell surface GLUT4). This makes sense since the amount of GLUT4 transported to the cell surface fluctuates as the energy needs of the cell vary.

## Discussion

In this paper, we have developed a framework and a corresponding computational package which analytically derive the steady states of a large class of chemical reaction networks. This framework utilizes network decomposition to break a CRN into smaller and more manageable pieces. This is significantly more efficient than the previous methods of solving steady states presented in [20, 21], which required performing the network translation on whole networks that are not WR and DZ, which could be very challenging if the given network is large and complicated. Using our approach, we can derive the steady state of a CRN with the mass-action kinetics if each independent subnetwork is either a WR and DZ network, or can be transformed into a WR and DZ network. To facilitate this, we developed a user-friendly and publicly available computational package, called COMPILES.

Previously, enormous number of numerical simulations were performed to investigate the bistability of the CRISPRi toggle switch model (Fig. 3) even within a limited range of parameters [27]. However, in this work, the closed form of its steady state, derived with our approach, allows us to easily confirm the absence of bistablility. Specifically, the closed form allows the flexibility of investigating the steady state concentrations of various species with respect to varying initial concentrations and parameters. This elucidates the monotonicity of the steady state concentration of S (single guide RNA 1) over varying the rate constants, which confirms that the model could not produce bistability in species *S*. Hence, this predicts that additional reactions had to be introduced to the network to capture the experimentally observed bistability in the system [27].

Furthermore, we were able to easily obtain the analytic steady state of the insulin model with the help of COMPILES, which allows us to quickly detect which species have absolute concentration robustness (ACR). That is, our method identifies those species whose steady state concentrations are maintained within a narrow range, regardless of the changes in the amounts of other species in the network [14].

Solving steady states numerically is common for establishing multistability, performing sensitivity analysis, conducting bifurcation analysis, and determining steady state stability. Numerical approaches, however, typically involve enormous amount of computation and investigate a limited range of parameter values. Solving these states analytically, on the other hand, allows these procedures to be done efficiently and within a wider range of parameters.

While we focus on the steady states of deterministic systems, the stationary distributions of stochastic systems can also be derived analytically for WR and DZ biochemical reaction networks [22, 24, 33, 34]. It would be interesting in future work to investigate whether the combination of network decomposition and translation can be used to derive stationary distributions analytically for a large class of biochemical reaction networks.

## Methods

### Chemical reaction networks

A *chemical reaction network* (CRN) can be seen as a finite collection of unique *reactions*. For instance, the following network, denoted by 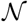, is a CRN with four reactions:

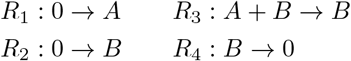

and fundamental units *A* and *B* called *species*.

In this CRN, reactions *R*_1_ and *R*_2_ indicate the production of species *A* and *B*, respectively. Additionally, *R*_3_ signifies that the encounter between *A* and *B* results in the disappearance of *A*. Finally, *R*_4_ designates the consumption of *B*.

The structure of a CRN can be easily viewed as a directed graph where the edges are the reactions, and the *nodes* are non-negative linear combinations of the species *A* and *B* in the network. In particular, *R*_3_ : *A* + *B* → *B* has nodes *A* + *B* and *B*, which are called the *source* node (before the arrow) and the *product* node (after the arrow), respectively. Hence, we are in a position to say that a CRN is composed of these three sets: the sets of species, nodes (also called *complexes*), and reactions [7, 26].

We associate each reaction of a CRN with the difference between its product and source nodes called a *reaction vector*. Thus, the reaction vectors for *R*_1_, *R*_2_, *R*_3_, and *R*_4_ are given as follows:

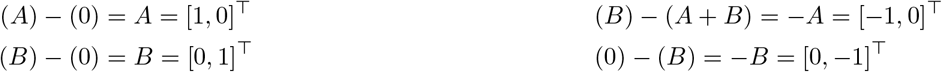

by identifying the two species *A* and *B* with the standard basis column vectors [1, 0]^⊤^ and [0,1]^⊤^ of the Euclidean space ℝ^2^, respectively. Then, the *stoichiometric matrix* of 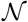 is a matrix whose columns are the reaction vectors, and is written as

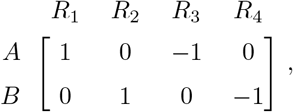

whose rank is given by *s* = 2.

With the rank of this matrix, we can define a very important concept in CRN theory called the *deficiency* of a CRN [8–10], denoted by *δ*, which can be easily calculated using the formula *δ* = *n* – ℓ – *s*, where *n* is the number of nodes, ℓ is the number of connected components, and *s* is the rank of the stoichiometric matrix. The deficiency of a CRN is a non-negative integer that can be interpreted as the measure of linear independence among its reactions. Networks with *δ* = 0 have the highest linear independence among the reactions. On the other hand, a high value of *δ* of a CRN means that linear independence is low [15].

A CRN is usually endowed with a kinetics to describe the evolution of the concentration of the species over time, which forms the *chemical reaction system*. Specifically, when the kinetics is *mass-action*, the rate function of each reaction is proportional to the the product of the concentration of the species in its source node. Hence, this rate function is a proportionality constant (*rate constant*) times the product of each concentration raised to the stoichiometric coefficient of the species that occurs in the source node of the associated reaction.

Suppose that *a* and *b* are the concentrations of the species *A* and *B*, which evolve over time. Then, the rate functions are given as follows:

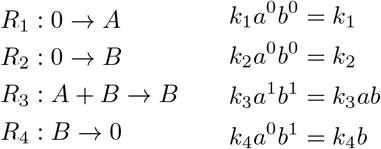

where *k_i_* is the rate constant of the reaction *R_i_*.

Thus, the set of ordinary differential equations (ODEs) that describes the dynamics of the mass action system is the following:

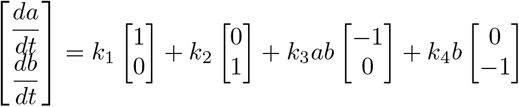

where 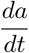 and 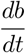 are the time derivatives of the concentration functions of the species *A* and *B*, respectively. Furthermore, the equation can be written as

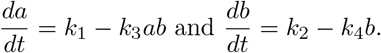

The long-term behavior of a system is described by *steady states*, which could be solved by equating each time derivative to zero, i.e., 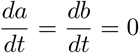, and solving for the concentration of the species in terms of the rate constants. Hence, *k*_1_ – *k*_3_*ab* = 0 and *k*_2_ – *k*_4_*b* = 0, so the steady state is 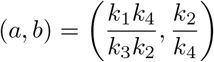 where *k*_1_, *k*_2_, *k*_3_, *k*_4_ > 0.

Alternatively, the steady state can be observed from a simulation of the ODEs but for particular rate constants of the reactions and specific initial values of the concentrations of the species. Although, in general this approach is used, especially when the network is large and complicated, because it is easier to simulate rather than derive the closed form of the steady state, important properties of steady states such as existence and uniqueness are difficult to justify using numerical simulations.

### Network decomposition

A *decomposition* of a CRN is induced by a partition of its reaction set. Suppose that we partition the reaction set 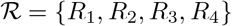 of the CRN 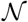 into the following subsets:

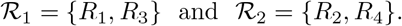

This partition then induces a decomposition of the network into two subnetworks, which we label 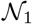 and 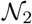.

In the case where the rank of stoichiometric matrix of the whole network is the sum of the ranks of the stoichiometric matrices of its subnetworks, then decomposition is called *independent* and we refer to the subnetworks as *independent subnetworks* [25, 26]. Specifically, the stoichiometric matrices of 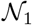 and 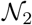 are

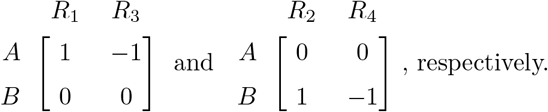

The ranks of these matrices are both one, i.e., *s*_1_ = *s*_2_ = 1. Since the rank of the entire network is *s* = 2 = 1 + 1 = *s*_1_ + *s*_2_, then the decomposition is independent. In general, to obtain the finest independent decomposition (the independent decomposition with the maximum number of subnetworks), we follow the method introduced in [35, 36] (see the Supplementary Information for details).

It was shown that when the underlying network decomposition of a reaction network is independent, then the set of steady states of the whole system is equal to the intersection of the sets of steady states of the subsystems as long as the same kinetics is followed by each reaction from the whole network down to its corresponding subnetwork [25, 26] (see Supplementary Information for details).

### Computational package, COMPILES

We developed a user-friendly, open-source, and publicly available computational package, COMPILES, that automatically decomposes a CRN into its finest independent decomposition. The package then derives the steady state of each subnetwork using the method outlined in Figs. 1b and 2. Finally, COMPILES combines these subnetwork solutions to output the analytic steady state solution for the entire network in terms of the rate constants and free parameters. It also gives additional information by enumerating the conservation laws of the system.

To efficiently solve each subnetwork, a subnetwork that is already weakly reversible and of deficiency zero is no longer translated. COMPILES does its best to output the most simplified analytic solution in terms of rate constants and free parameters. If this is not feasible, the unsimplified solution (with non-free parameters) is returned.

## Supporting information

**S1 Appendix. Technical details of the study.** This file contains the theoretical aspect of the study and the technical details of the main examples and the computational package.

## Acknowledgments

BSH and JKK are supported by the Institute for Basic Science IBS-R029-C3. MDJ is supported by NSF grant DMS-2213390. The authors acknowledge Hyukpyo Hong and Yunmin Song for helpful discussions.

## Author contributions

**Conceptualization:** Bryan S. Hernandez, Patrick Vincent N. Lubenia, Matthew D. Johnston, Jae Kyoung Kim.

**Funding acquisition:** Jae Kyoung Kim.

**Methodology:** Bryan S. Hernandez, Patrick Vincent N. Lubenia, Matthew D. Johnston, Jae Kyoung Kim.

**Project administration:** Jae Kyoung Kim.

**Software:** Patrick Vincent N. Lubenia.

**Supervision:** Jae Kyoung Kim.

**Visualization:** Bryan S. Hernandez, Jae Kyoung Kim.

**Writing – original draft:** Bryan S. Hernandez, Patrick Vincent N. Lubenia, Matthew D. Johnston, Jae Kyoung Kim.

**Writing – review & editing:** Bryan S. Hernandez, Matthew D. Johnston, Jae Kyoung Kim.

